# Energy efficiency and sensitivity benefits in a motion processing adaptive recurrent neural network

**DOI:** 10.1101/2024.12.20.629845

**Authors:** Vishnu Mohan, Reuben Rideaux

## Abstract

Motion processing is a key function for the survival of many organisms and is initially implemented in the primary visual cortex (V1) and the middle temporal area (V5/MT) of the primate visual cortex. Advances in machine learning approaches have led to the development of motion processing neural networks that have elucidated several aspects of this process. However, it remains unclear how adaptation, a canonical function of sensory processing, influences motion processing. In this study, we developed two recurrent neural networks to study motion processing: MotionNet-R, a baseline model, and AdaptNet, a model that employs adaptive mechanisms inspired by biological systems. Both networks were trained on natural image sequences to estimate motion vectors. We found that both networks developed response properties that resembled those of neurons found in areas V1 and MT, e.g., speed tuning, and AdaptNet recapitulated the motion aftereffect phenomenon (i.e., *the waterfall illusion*). We show that the emergent computational properties that implement the phenomenon in AdaptNet confirm previous theoretical hypotheses. Further, we compared the performance of the two networks and found that AdaptNet processed motion more efficiently, operationalized as reduced activation. While AdaptNet incurred reduced accuracy in response to prolonged constant input, it was both more accurate and sensitive in response to changes in motion input. These results are consistent with theoretical explanations of adaptation as neural property that supports metabolic efficiency and increased sensitivity to change in the environment. Our findings provide novel insights into the neural mechanisms underlying motion adaptation and highlight the potential advantages of adaptive neural networks in modelling biological processes.

## INTRODUCTION

### Biological Basis of Motion Processing and Adaptation

The ability to process and interpret motion is one of the most fundamental requirements for survival in any environment. In the primate visual cortex, multiple areas work together in a hierarchical fashion to process increasingly complex forms of motion. Two key regions in this network are the primary visual cortex (V1) and the middle temporal area (V5/MT), which serve intrinsic roles in the early stages of motion processing. While V1 is primarily responsible for detecting local motion, MT integrates these local signals to compute global motion (Movshon & Newsome, 1996). This processing hierarchy in perhaps most prominently demonstrated by the responses to V1 and MT neurons in response to drifting sinewave and plaid gratings (Movshon et al., 1985).

A key feature of motion processing, and healthy brain function more broadly, is adaptation. Motion adaptation manifests in various perceptual phenomena, most notably the motion aftereffect (MAE) or “waterfall illusion”, where prolonged exposure to motion in one direction leads to perception of motion in the opposite direction when subsequently viewing a stationary stimulus (Mather et al., 1998). While the MAE itself does not seem to enhance perception, it reflects underlying adaptive mechanisms thought to optimize neural resources and enhance sensitivity to changes in visual motion (Mather et al., 1998).

There have been significant advances in our understanding of motion processing (Born & Bradley, 2005; Burr & Thompson, 2011; Nishida, 2011) and its limitations (Edwards & Rideaux, 2013; Rideaux & Edwards, 2014). However, the neural mechanisms that underlie motion adaptation are not yet fully understood (Clifford et al., 2007; Glasser et al., 2011; Solomon & Kohn, 2014). Multiple factors contribute to this challenge. One such issue is that motion adaptation is distributed across several areas of the brain, which makes it difficult to isolate its effects on specific areas (Britten, 2008). Even within a single brain region, there is significant variability in both anatomy and response properties. For example in V1, the diversity of cell types, subcortical input adaptation, and complex changes in both gain and tuning preference of neurons present additional challenges (Carandini et al., 1999; Wissig & Kohn, 2012). This is further complicated by the wide range of adaptation timescales, from milliseconds to hours, which makes comprehensive experimental design difficult (Glasser et al., 2011). There is also an inherent variability in neural responses, which obscures adaptive changes (Bair & Movshon, 2004). Studying adaptation in higher layers poses additional challenges due to the interplay between inherited and local adaptation effects, which can be difficult to disentangle (Patterson et al., 2013; Rideaux & Harrison, 2019; Zamboni et al., 2020). Additionally, factors such as the diversity of cell types in higher regions like area MT, which adapt differently to those in V1 (Ll et al., 2007), and the potential for top-down influences such as attention (Kohn & Movshon, 2004) further complicate the study of motion adaptation.

Despite of the difficulty associated with studying motion adaptation, considerable progress has been made towards understanding this phenomenon. Johnston et al. (2006) revealed the effect of adaptation on speed perception, while the influence of adaptation on motion integration was explored by Patterson et al. (2014). Lee (2018) further built upon this by exploring the interplay between local and global motion under adaptation, uncovering bidirectional effects. Notably, adaptation can also improve motion detection capabilities in noisy conditions by improving sensitivity to motion signals that are relatively weaker (Hietanen et al., 2007). The scope of motion adaptation extends beyond simple 2D motion; for example, Sakano and Allison (2014) investigated motion-in-depth perception and found that adaptation to motion towards or away from an observer can induce an aftereffect in the opposite direction. Adaptation also impacts higher-level processes, such as biological motion perception in humans, as demonstrated by Theusner et al. (2011), who showed that adaptation to a walking figure can bias the perceived walking direction of a subsequently viewed figure. Additionally, Cuturi and MacNeilage (2014) highlighted that adaptation affects vestibular heading perception, illustrating its role in multisensory integration. This diverse body of research demonstrates how pervasive the effects of adaptation are across different forms of motion perception.

### Computational Approaches to Motion Perception

Computational modelling has shown to be an effective tool for studying motion processing. Over the last few decades, there have been pioneering computational models of visual processing that have focused on both ventral and dorsal streams. In particular, Simoncelli and Heeger (1998) proposed an influential model of MT neurons that accounted for their direction and speed tuning. Rust et al., (2006) expanded on this work by implementing a more detailed and biologically plausible V1 layer. These early models laid the groundwork to study motion processing in area MT using artificial networks.

In recent years, computational modeling combined with machine learning has increasingly been applied to investigate complex biological phenomena (Kriegeskorte & Douglas, 2018). Deep neural networks have been a useful tool for understanding the hierarchical structure of processing in the brain (Yamins & DiCarlo, 2016). Training these networks on natural images leads to representations that mimic those observed in the primate visual cortex (Cadieu et al., 2014; Guclu & Van Gerven, 2015; Khaligh-Razavi & Kriegeskorte, 2014). While many studies have used deep neural networks to investigate visual processing, they have predominantly focused on ventral stream functionality. Notable examples include the work of Yamins et al. (2014), who used deep learning to show that trained networks develop internal representations like those observed in higher visual areas of the ventral stream, and Kar et al. (2019), who used convolutional neural networks (CNNs) to predict responses in the ventral pathway during object recognition tasks.

There have been some more recent efforts to utilize machine learning techniques to understand dorsal stream processes. For instance, Rideaux and Welchman (2020) used a CNN trained on natural image sequences to model motion processing, successfully reproducing key properties of V1 and MT neurons and providing novel insights into the origins of motion illusions. Further, Qiao and Shen (2021) developed a model that learned to estimate motion from natural human videos and Gundavarapu et al. (2019) proposed a hierarchical model of motion processing that learned representations like those in the dorsal pathway. However, while the above artificial neural network (ANN) models successfully reproduced certain qualities of processing in the visual cortex, the effects of adaptive mechanisms on those qualities have been neglected.

Parallel to these developments, there has been considerable work investigating the benefits of adaptation in the field of neuromorphic computing. For example, adaptive mechanisms in spiking neural networks (SNN) produce efficiency gains (Ganguly et al., 2024; Indiveri & Liu, 2015) and optimizes sensory encoding (Mao et al., 2025) and the use of neuron models such as adaptive exponential neurons (AdEx) along with neuromorphic hardware has led to considerable gains in energy efficiency for certain tasks (Bellec et al., 2018, 2020). Such mechanisms have also been shown to enhance performanve in situations involving a change detection (Hu et al., 2021). These studies highlight the importance of adaptation in spiking neural networks and suggest that incorporating adaptation can enhance computational efficiency in neuromorphic systems. However, the influence of adaptation in ANNs trained to perform visual tasks, in particular visual motion estimation, remains unexplored.

### Present Work

While computational models of motion processing have provided considerable insights into this process, our understanding remains incomplete. Incorporating adaptation into neural networks may expand our understanding of motion processing across different areas of the visual cortex. Thus, here we developed novel ANNs that simulate motion processing and adaptation in the visual cortex, focusing on early to mid-level visual areas V1 and MT. These ANNs incorporate adaptation mechanisms in both V1- and MT-like layers, allowing us to study the role of adaptation in motion processing. By comparing the emergent properties of adaptive and non-adaptive networks trained on natural image sequences, we reveal novel insights into the neural mechanisms underlying motion adaptation in visual processing, complementing and extending previous theoretical and empirical work.

## METHODS

Two convolutional recurrent neural networks (conv-RNNs) were trained to estimate motion velocity from natural image sequences: MotionNet-R, a baseline model, and AdaptNet, which employs adaptive mechanisms. Our goal was to better understand the role of adaptation in motion processing by comparing the emergent properties within these networks. This comparison allows us to directly measure how adaptation mechanisms can influence motion processing, within a neural network.

### Training data

The network was trained with 60,000 natural image sequences, each containing ten frames, extracted from photographs in the Berkeley Segmentation Dataset (Martin et al., 2001). This approach has previously been used (Rideaux & Welchman, 2020, 2021; Simoncelli & Olshausen, 2001) and aims to reproduce the visual diet of biological visual systems. A sliding window method was used to generate motion sequences from natural images. The ground truth velocity values corresponding to each sequence were recorded during this process, for use in training. The sequences were kept relatively short to mitigate vanishing gradients.

### Model architecture and theoretical foundations

We developed two ANNs to simulate motion processing and adaptation in the visual cortex, focusing on early to mid-level visual areas V1 and MT. The first network, which we named MotionNet-R, expands upon the original MotionNet architecture (Rideaux & Welchman, 2020, 2021) by incorporating a recurrent layer. The second network, AdaptNet, further builds on MotionNet-R by implementing adaptation mechanisms in both V1 and MT units. The only difference between MotionNet-R and AdaptNet was the implementation of the adaptation mechanism in the convolutional and recurrent layers. The structure of the two networks was the same: a two-dimensional (2D) convolutional layer (simulating V1), a recurrent layer (simulating MT), and a linear output layer. The networks’ structure was intended to recapitulate the hierarchical organization of areas in the visual cortex (Born & Bradley, 2005; Movshon & Newsome, 1996). The fine tuning of parameters for training was done by trial and error. A dropout layer was added after the convolutional and recurrent layers to improve regularization (Srivastava et al., 2014), and make the training more robust. The dropout did not interact directly with the adaptation mechanism.

### Convolutional layer (V1 simulation)

Following the input layer, the first hidden layer of the networks was a 2D convolutional layer with 16 kernels (size = 6×6 pixels; stride length = 1). This 2D convolutional layer has 2 input channels corresponding to consecutive frames of the input. These values were selected based on previous work (Rideaux & Welchman, 2020) to minimize network scale and complexity. Two successive images of the motion sequence were fed as input to this layer, with each image serving as a separate channel. The operation of a 2D convolutional layer over [i, j] with 2 input channels can be described as:

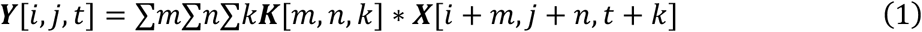

where ***Y*** is the output, ***X*** is the input, ***K*** is the kernel, *i*, *j* are spatial coordinates, *m*, *n*, *k* are the kernel dimensions and * denotes convolution.

### Recurrent neural network layer (MT simulation)

Area MT was modelled using a recurrent layer in both networks. This layer was responsible for analyzing temporal information across a series of V1 responses. The operation of area MT and other areas in the neocortex have previously been modelled with recurrent networks (Kar et al., 2019). Each unit in this layer uses a nonlinear activation function (rectified linear unit; ReLU) to introduce nonlinearity into the network. The ReLU function is defined as:

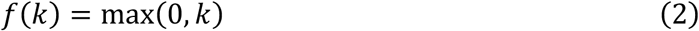

where *k* is the input activation at a ReLU unit.

### Adaptation mechanism

In AdaptNet, we implemented an adaptation mechanism in both V1 and MT units. We aimed to model this adaptive mechanism based on what is observed in biological neural networks like the visual area of the macaque (Kohn & Movshon, 2003; Krekelberg et al., 2006). Key to this implementation was an adaptation variable that accumulates over time (to reflect a decrease in response to a constant stimulus) and gradually decays (recovery). This adaptation was described mathematically as:

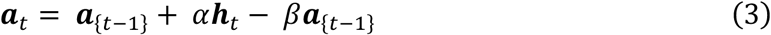

where ***a****_t_* is the adaptation at time *t*, *h_t_* is the hidden state (activation) at time *t*, *α* is the adaptation rate, and *β* is the recovery rate. The adaptation variable accumulates when the neuron is active (controlled by ***a****_t_*) and decays over time (controlled by recovery rate *β*). A higher adaptation rate (***a****_t_*) in means these units show stronger or faster response reduction with sustained firing. We used an adaptation rate of 0.2 for the MT units and 0.1, for the V1 units. A recovery rate of 0.1 was used in both cases. In MotionNet-R, we omit the adaptation variable and associated adaptive logic. Hence, the recurrent layer and convolution layer outputs are passed as is, to the subsequent layer. We used a lower adaptation rate in the V1 layer compared to the MT layer to reflect the same difference in adaptation observed in V1 and MT of biological neural networks like those of the macaque (Patterson et al., 2014; Priebe et al., 2002; Solomon & Kohn, 2014).

### Linear layer (readout)

The output layer of both networks was a linear layer that mapped the activity of the recurrent layer to two outputs: x and y. Here, x and y denote the estimated motion velocity (in pixels per frame) along the horizontal and vertical axes, respectively. Utilization of area MT’s responses to estimate speed and direction has been observed in humans and adult male rhesus monkeys (Churchland & Lisberger, 2001). The operation of this layer can be described as:

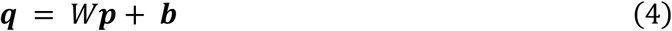

where ***q*** is the output (motion estimate), ***p*** is the input from the recurrent layer, ***W*** is a weight matrix, and *b* is a bias vector.

### Training procedure and evaluation

Both networks were trained using mean squared error (MSE) loss and the Adam optimizer (Kingma & Ba, 2017). The MSE is defined as:

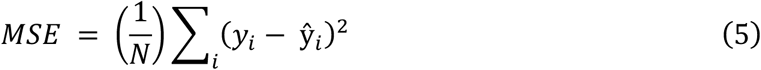

where *N* is the number of samples in a batch of data, *y_i_* are the true values and 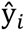 are the predicted values.

We trained ten separate instances of the networks, each for 30 epochs. The variability between the models stemmed from random initializations and varied order of minibatch sampling. Having multiple networks to test allows us to quantify the variability of the results (Kriegeskorte & Douglas, 2018). The difference between the network’s predictions and ground truth was quantified using Pearson’s correlation coefficient.

### Spatial and temporal frequency tuning analysis

We extracted response profiles for each unit by exposing the network to drifting sinewave gratings and moving dot sequences, across a range of spatial and temporal frequencies (10 frame sequences, 32×32 pixels, moving dot diameter was 3 pixels unless specified otherwise). This approach mimics that used in neurophysiological experiments to characterize the tuning functions of neurons (De Valois et al., 1982). We generated sinewave grating stimuli using the following equation:

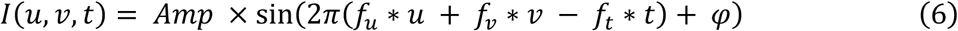

where *I*(*u*, *v*, *t*) is the intensity at position (*u*, *v*) and time *t*, *Amp* is the amplitude (contrast), *f_u_* and *f_v_* are the spatial frequencies in the *u* and *v* directions, *f_t_* is the temporal frequency, and *φ* is the phase.

We also tested the networks on plaid stimuli (generated by combining two sinewave gratings) to evaluate the units’ responses to component and pattern motion. We varied spatial frequency logarithmically from 0.01 to 1 cycles/pixel and temporal frequency from 0.1 to 5 Hz. For each combination of spatial and temporal frequency, we presented gratings moving in eight different directions. For some of our analyses, directional and speed preferences were similarly calculated using moving dot sequences that varied in speed within a range between ±3 pixels/frame. The preferred direction, spatial and temporal frequencies of each V1 and MT unit was determined by identifying each unit’s maximal response to the varying stimuli.

### Visualization of convolutional kernels

To gain insight into the features detected by the convolutional layer, we visualized the trained convolutional kernels (Zeiler & Fergus, 2013). We normalized the weights of each convolutional kernel ***K*** using the following equation:

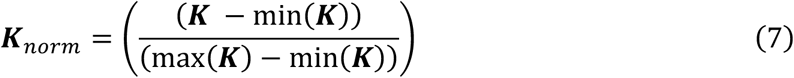

We then applied Gaussian smoothing to highlight edge patterns, using the equation

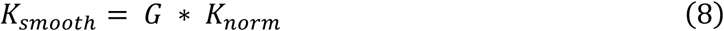

where *K_smooth_* is the smoothed kernel, *K_norm_* is the normalized kernel, * denoted 2D convolution and ***G*** is a 2D Gaussian kernel defined as

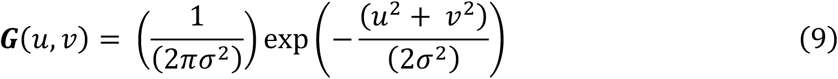

where *u*, *v* are the spatial coordinates in the 2D Gaussian kernel and *σ* is the standard deviation that controles the amount of smoothing. We used σ = 1 for smoothing.

### Analysis of weight connections between directionally tuned units

To investigate how information is processed by the networks to support motion estimation, we analyzed the connections between V1 and MT units (V1-to-MT), and the recurrent connections between MT units (MT-to-MT) (Qian & Andersen, 1994; Snowden et al., 1992). We computed each V1 unit’s average V1-to-MT weight by first aligning its preferred direction, and those of the MT units, to zero, before binning and averaging all weights within eight linearly spaced directions from 0-360 degrees.

### Waterfall effect analysis

To test for perceptual phenomena associated with adaptation in AdaptNet, we investigated whether the network exhibited the waterfall illusion, a well-known motion aftereffect (Mather et al., 1998). We modified testing sequences to probe for an aftereffect (natural image sequences that were 11 frames long; stationary initially for two frames to gauge any inherent network bias, then three frames of movement followed by six frames of stationary stimulus), presented this sequence to AdaptNet, and analyzed the output.

### Efficiency analysis

We tested the efficiency of both networks by comparing their response magnitude and accuracy. We exposed both networks to two types of moving dot sequences; one where the speed and/or direction of the moving dot varied every five frames and another kind where the moving dot maintained a constant trajectory with the same speed, throughout the duration of exposure. The MSE was calculated for a window of three frames after each input change point. Recurrent unit outputs across all frames were then summed to estimate the energy used by the network to sustain a representation. Our aim was to test if AdaptNet displayed increased efficiency compared to MotionNet-R. Such an increase in efficiency with adaptation is supported by studies of biological visual systems (Barlow & Földiàgk, 1989; Wainwright, 1999). Practically, the test would help indicate if the reduced overall activation displayed by AdaptNet would lead to a lower net output when compared to MotionNet-R, without a substantial relative decrease in accuracy.

### Sensitivity Analysis

To examine the difference in sensitivity to a varying input stimulus between the networks, we exposed them to a variety of moving dot sequences with varied speeds. The sequences consisted of ten image frames, of which the first nine frames were one speed and the tenth frame was either the same or a different speed (32×32 images, speed varied from ±3 pixels/frame). This led to several initial-final speed combinations; for each of these combinations we presented 100 image sequences, each with varied initial dot locations. For each combination, we measured the networks’ accuracy (as MSE) in estimating the correct motion direction of the final speed (separately) along x and y dimensions. This analysis produced a probability matrix that we could then compare with the ground truth. We then fit a sigmoid function to the diagonal values of these matrices to estimate the sensitivity of each network to stimuli with no change in final speed (congruent condition). The non-diagonal elements were averaged across the range of initial speeds and a sigmoid was fit to this data to estimate each networks’ sensitivity to stimuli where final speeds varied from the initial speed (incongruent condition).

### Data availability

The code used to train and test the networks is publicly available at https://github.com/Vishnu-Mohan-USyd/Energy_efficiency_and_sensitivity_benefits_in_a_motion_processing_adaptive_recurrent_neural_network.git

## RESULTS

### Network performance evolution during training

Both MotionNet-R and AdaptNet learned to estimate motion reliably, following training with natural images. Both networks were structured with a convolutional layer simulating V1, a recurrent layer simulating MT, and a linear output layer for motion estimation (**Fig. 1a**). MotionNet-R displayed increasing accuracy over successive input frames (**Fig. 1b**), demonstrating the network’s ability to integrate information over time to support reliable motion estimation. By contrast, while AdaptNet’s accuracy initially increased with successive frames, it then subsequently decreased with further frames (**Fig. 1c**). This pattern reflects the impact of adaptation on the network’s performance. In particular, the initial increase in accuracy is likely driven by temporal integration, but then the adaptation induced by the constant motion causes the network’s response magnitude to decrease and the estimate to deviate from the ground truth, thus reducing the accuracy of motion estimation in response to later frames.

**Figure 1.**
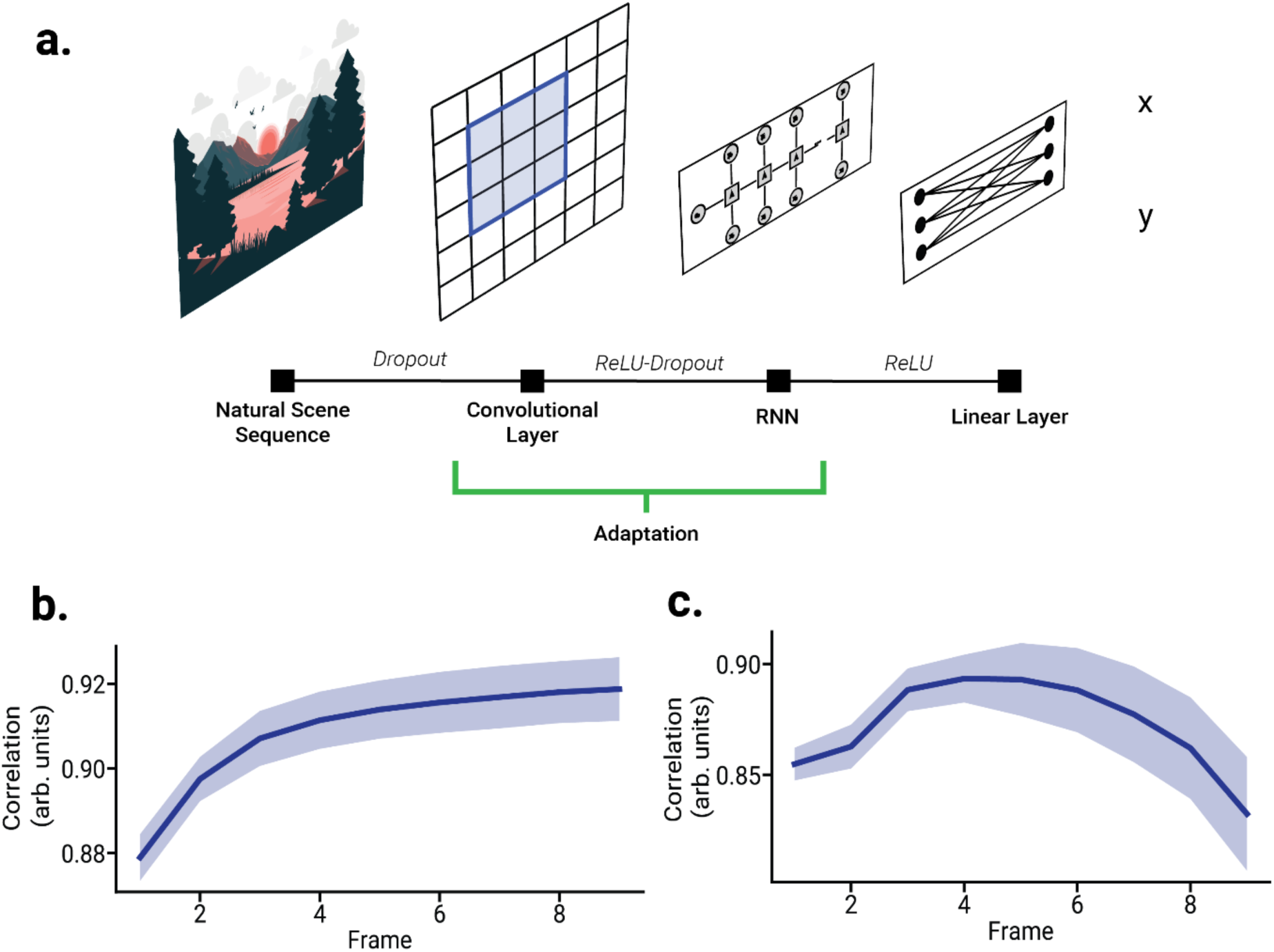
Network Architecture and Performance. **a**) Schematic diagram of the network architecture, showing the convolutional layer (simulating V1), the recurrent layer (simulating MT), and the linear output layer for motion estimation. **b, c**) Performance of (**b**) MotionNet-R and (**c**) AdaptNet, showing the correlation coefficient between predicted and ground truth motion values over time (successive image sequence frames). Shaded regions indicate ±SD across multiple networks.

### Spatiotemporal tuning properties in the networks

Both networks developed response properties resembling those of biological neurons in areas V1 and MT. The distribution of speed tuning among convolutional (V1) and recurrent (MT) units was similar to neurophysiological observations in macaque (Maunsell & Van Essen, 1983; Priebe et al., 2006) (**Fig. 2a-b**). That is, V1 units were tuned to relatively lower speeds than MT units. This resembles the responses of neurons in area V1, which are typically more sensitive to lower velocities when compared to those of area MT.

**Figure 2.**
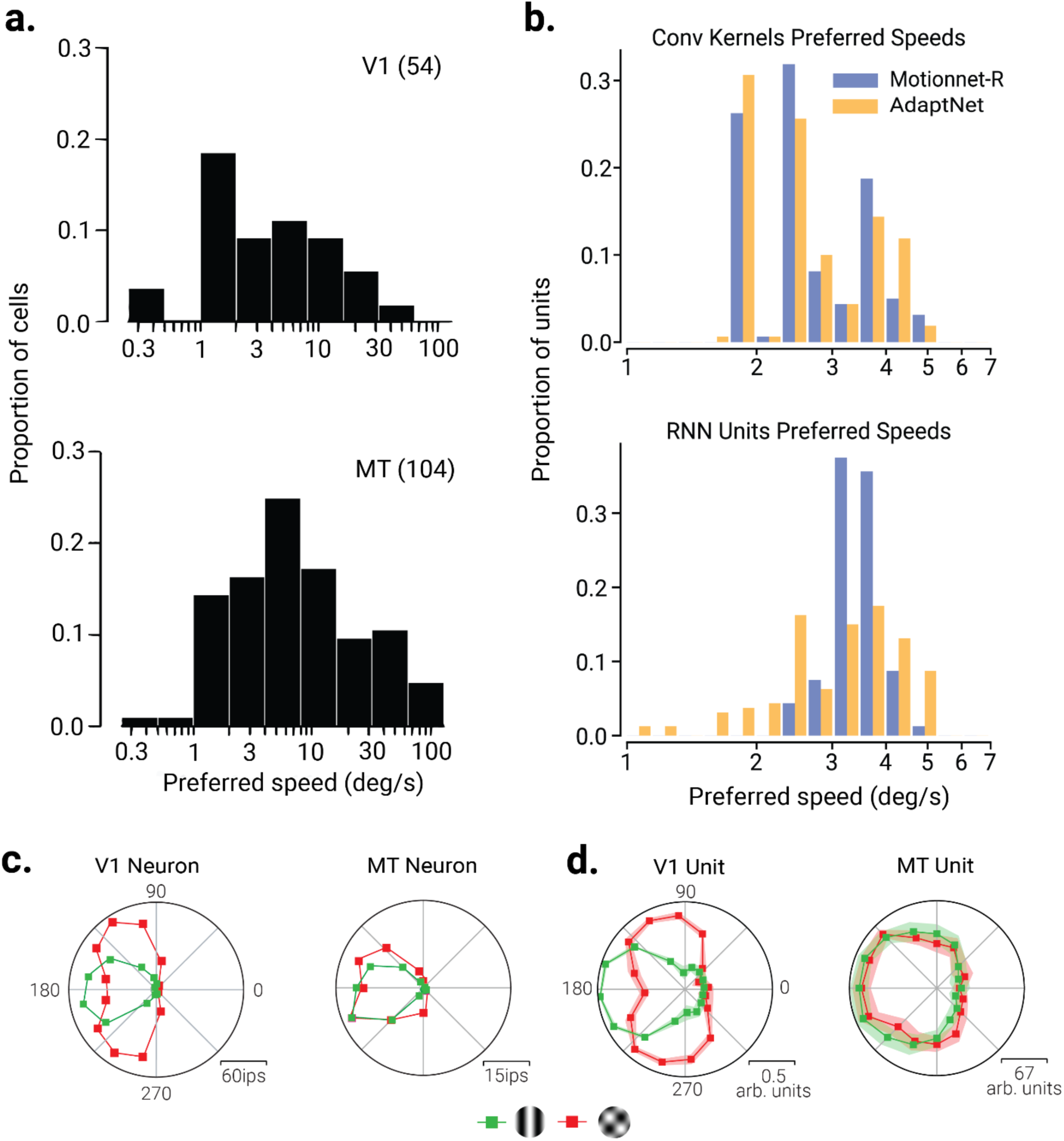
Spatiotemporal tuning properties of network units. **a)** Data from Priebe, et al. (2006) showing the distribution of preferred speeds for (top) V1 and (bottom) MT neurons in macaque. **b)** The same as (a), but for V1 and MT units in MotionNet-R and AdaptNet networks. **c)** Replotted data from Adelson and Movshon (1982) showing single-neuron responses in (left) V1 and (right) MT to sinewave grating and plaid stimuli (135° separation) moving in different directions. **d)** Same as (c), but for responses obtained from MotionNet-R units. Shaded regions indicate ±SD across multiple networks.

A key distinction between V1 and MT neurons is that the former responds maximally to component motion, while the latter responds to pattern motion, which is thought to be due to pooling of signals between V1 and MT. This phenomenon is clearly observable in the responses of these neurons to drifting sinewave grating and plaid stimuli. V1 neurons respond maximally to the two components of the plaid stimuli, whereas the MT neurons maximally respond to the combined pattern motion, which is consistent with perception of these stimuli (Movshon et al., 1985; Rust et al., 2006) (**Fig. 2c**). We tested the networks with these stimuli and found the same pattern of results; V1 units were selective for component motion while MT units responded maximally to the pattern motion (**Fig. 2d**). Together, the spatiotemporal tuning properties that emerged in the networks to perform motion estimation appear to resemble those observed in biological systems, e.g., non-human primates. The emergence of pattern motion selectivity in the higher layers occurs even in simpler models without recurrence or adaptation (Rideaux & Welchman, 2020) as it simply relies on the pooling of component motion signals from earlier layers.

### Emergent weight properties in the networks

We analyzed the weights of MotionNet-R and AdaptNet to understand how the connectivity patterns in these networks supported motion processing. **Figure 3a** shows the learned convolutional kernels, which resembled edge detectors or Gabor filters. A shift in the phase of these Gabor-like kernels between the two channels, which receive input from successive image sequence frames, appears to have emerged to facilitate detection of motion energy. These kernels are analogous to the receptive fields of V1 simple and complex cells, which are sensitive to specific orientations and motion directions (DeAngelis et al., 1999; Hubel & Wiesel, 1962).

**Figure 3.**
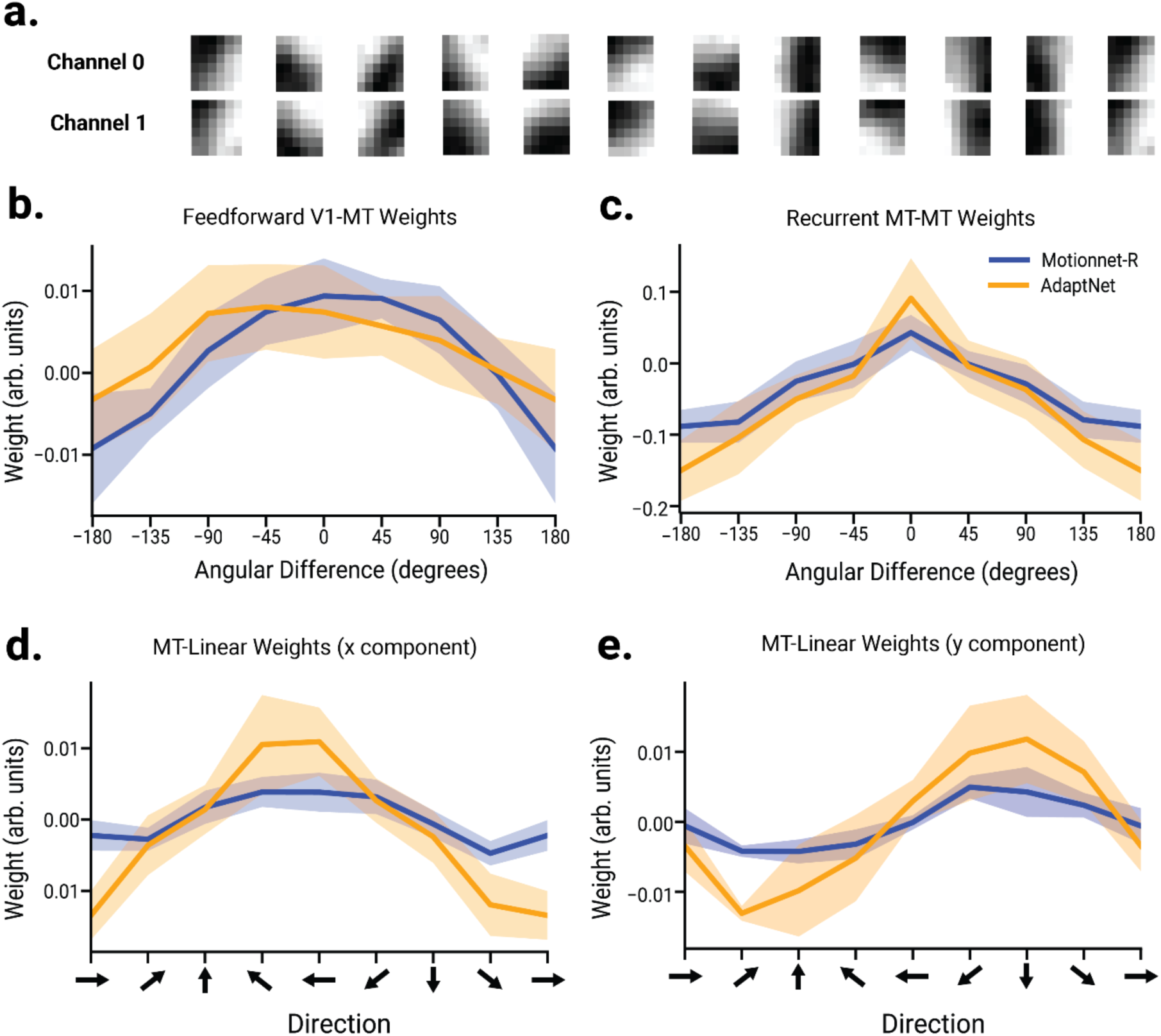
Network connectivity. **a**) Examples of convolutional (V1) kernels, resembling edge detectors or Gabor filters. Each column shows the two channels of a single kernel, corresponding to successive frames. **b**) Average feedforward connections from V1 to recurrent (MT) units as a function of directional tuning difference. **c**) Same as (**b**), but for recurrent connections between MT units (MT-to-MT). **d**, **e**) Average weights from MT units to the (**d**) x and (**e**) y component of the linear output layer as a function of the directional tuning of MT units. Shaded regions indicate ±SD across multiple networks. The legend in graph c also applies to graphs (b-e). Results are shown for both (blue) MotionNet-R and (orange) AdaptNet.

Assessment of the feedforward connections from V1 to MT units showed that there were more positive connections between units with a similar directional preference (**Fig. 3c**). This pattern of connectivity is consistent with previous neurophysiological observations between V1 and MT (Born & Bradley, 2005; Movshon & Newsome, 1996), and is thought to facilitate motion pooling (excitatory connections) and motion opponency (inhibitory connections). The recurrent connections between MT units (MT-to-MT) revealed that units with similar directional tuning are more positively connected (**Fig. 3b**). These connections likely serve similar motion pooling and noise reduction purposes as those between V1 and MT, and are analogous to lateral excitatory and suppressive synapses in the visual cortex (Li et al., 2014; Tan et al., 2011).

Recurrent connections were relatively balanced between excitation and inhibition (**Fig. 3c**), which is consistent with neurophysiology (for a review on the topic, see Zhou & Yu, 2018). In particular, the emergent connectivity was excitatory among units with similar motion direction preference and inhibitory among units tuned to dissimilar directions. Within the context of biological motion processing, this appears to be consistent with motion opponency mechanisms (Heeger et al., 1999)

**Figures 3d** and **3e** show the connections from recurrent units to the output layer responsible for estimating the x and y components of motion. As expected, we found that units tuned to horizontal movement contributed more to the x component output, while those tuned to vertical movement contributed more to the y component. While the V1-to-MT weights were stronger in MotionNet-R, the recurrent MT-to-MT connections were weaker. This may be a product of the increased adaptation in the MT units. Further, the increased connection strengths between the MT and output layer may reflect a compensatory effect within AdaptNet in response to the habituated (reduced) activity in the MT layer.

The emergent connections of the networks seem to mirror key motion processing aspects observed in biological visual areas V1 and MT. The alignment of the learned weights with neurophysiological connections in these areas supports the biological validity of the models.

### Adaptation effects on motion perception and network efficiency

Having found evidence supporting the biological validity of the emergent properties within the networks, we next tested how the networks respond to a stimulus that would typically elicit the motion aftereffect. The stimulus was comprised of three stages: an initial (baseline) stationary stage, an intermediate moving stage to induce adaptation, and then a final stationary stage to test for adaptation induced motion aftereffects (Fig. 4). As expected, MotionNet-R responded to the moving stimulus with increasing estimates, then gradually returned to baseline when the stimulus returned to stationary. By contrast, AdaptNet exhibited an overshoot in motion estimation following the cessation of stimulus movement, predicting motion in the opposite direction, which is a hallmark of the waterfall illusion observed in biological vision (Mather & Harris, 1998). To understand the computational basis for this motion aftereffect, we assessed the activity of the MT units across the different stages of stimulus processing (Fig. 4, top; the x output was studied in this case, the y output is similar, as shown in **Supplementary Fig. 1**). At first, before movement was initiated, MT units were responding equally in all directions, maintaining balanced baseline activity (Fig. 4, top-left). Once motion was initiated, units tuned to the motion direction increased their activity, while those tuned to other directions were suppressed (Fig. 4, top-middle). When the motion stopped, the adapted units recovered at different rates, leading to a temporary imbalance manifesting as the motion aftereffect (Fig. 4, top-right). The overshoot occurred for a range of input velocities (**Supplementary Fig. 2**). The mechanism underlying the motion aftereffect in the network aligns with previous hypotheses of the corresponding phenomenon in humans. Namely, that adaptation perturbs the balanced state of mutual suppression between motion-selective neurons, which leads to the illusory perception of motion in the opposite direction when the stimulus ceases (Grunewald & Lankheet, 1996; Kohn & Movshon, 2003). As can be seen in Figure 4, prolonged activation of a subpopulation of units preferring the stimulus motion direction leads to their activity being relatively lower than units tuned to other directions when the stimulus ceases to move.

**Figure 4.**
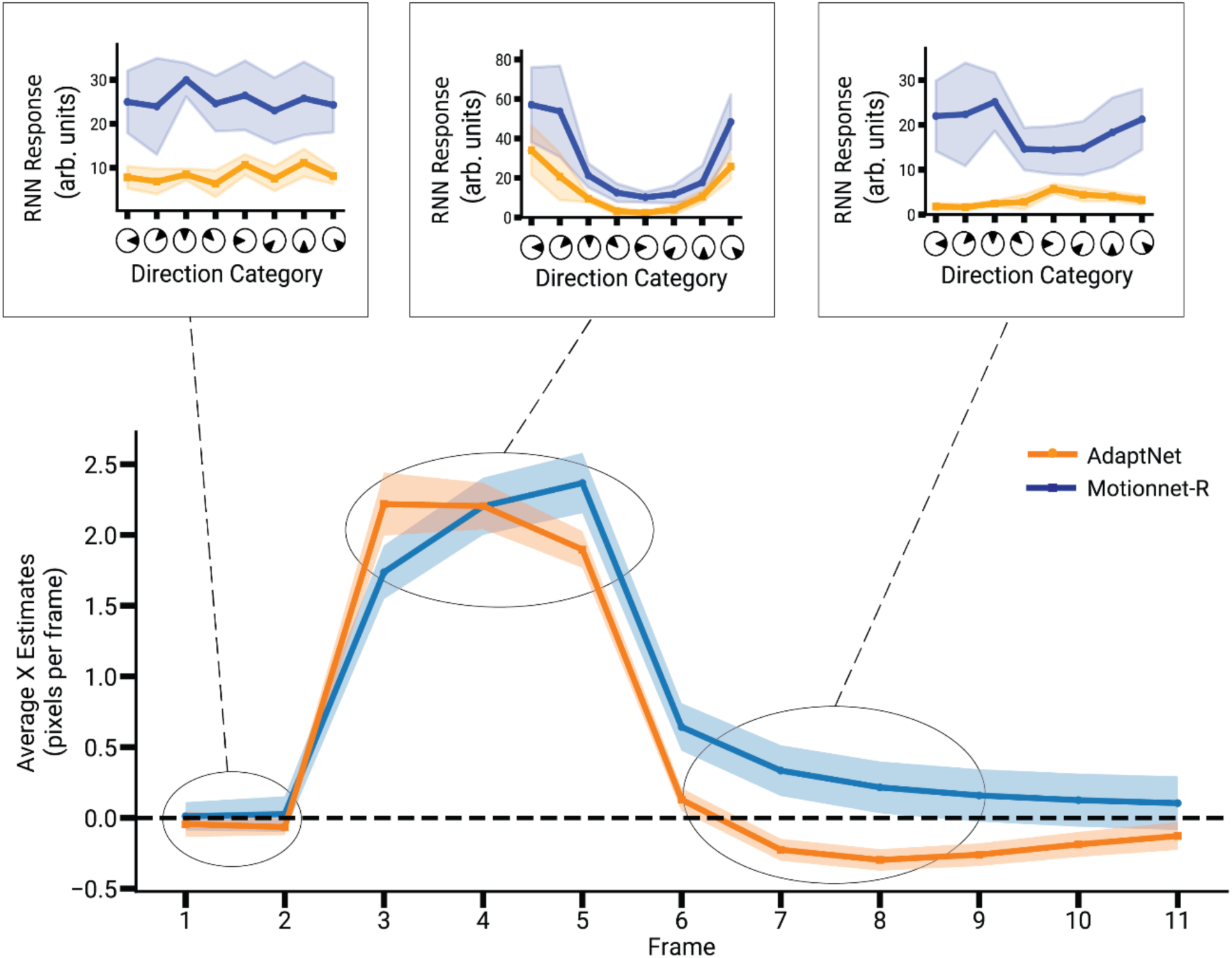
Motion Aftereffect in AdaptNet. Time course of the networks’ responses to an image sequence consisting of a stationary stage (2 frames), followed by a moving stage (3 frames), and then a return to stationary (5 frames). The lower plot shows MotionNet-R and AdaptNet’s motion estimation over time. The upper insets (showing RNN responses during frames 2, 4 and 7 representing the stationary, moving and final phases of the sequence respectively) show the average activity of MT units, binned according to their directional tuning, during different stages of the sequence. AdaptNet’s response moving into the opposite (negative) illustrates the aftereffect displayed by this network. Shaded regions indicate ±SD across multiple networks.

This overshoot was consistent across a range of input velocities, indicating that the adaptation mechanisms in the network can account for the perception of illusory motion under various conditions (**Supplementary Fig. 2**). This result aligns with previous hypotheses that motion aftereffects result from the adaptation of motion-sensitive neurons, causing shifts in the perceived direction of motion when the stimulus changes or ceases (Grunewald & Lankheet, 1996; Kohn & Movshon, 2003). In particular, we found that adaptation perturbed the homeostatic state of mutual suppression between RNN units when exposed to a stationary input. As can be seen in Figure 4a, prolonged activation of a subpopulation of neurons preferring one direction reduced their activity after the stimulus was removed (which is most apparen immediately after motion cessation, where AdaptNet’s response decreases to zero for the previously active units, while in MotionNet-R there is no such activation reduction). During this refractory period, the previously active neural group fails to provide an equal and opposing response as it would, during stationary stimulus presentation, and hence there is a temporary shift in prediction that favors the opposite direction group. This temporary overshoot in motion estimation gives rise to the perception of the illusory waterfall effect.

Another notable difference between the responses of the MotionNet-R and AdaptNet is the latency with which they respond to the initiation of motion. As can be seen in Figure 4, AdaptNet reaches peak velocity estimation more quickly than MotionNet-R. AdaptNet’s increased response speed may result from the reduced lateral suppression within the MT layer (passed from the previous time point). That is, the MT units tuned to the stimulus direction have less inertia to overcome at the onset of motion. To further investigate this phenomenon, we measured the average number of frames required by the MT units of each network to reach peak velocity estimation and found that AdaptNet consistently reached the peak 2-3 frames earlier than MotionNet-R, across a range of speeds (**Supplementary Fig. 3**).

Adaptation is thought to support healthy brain function by increasing sensitivity to changes in sensory input while reducing the otherwise high metabolic expenditure required for sustained cortical computation (Duong et al., 2023; Lennie, 2003). To test for this in our network, we compared the MSE between the actual and estimated motion, and the total MT layer activity, between AdaptNet and MotionNet-R in response to inputs with and without motion changes (Figure 5a). For image sequences of constant motion, AdaptNet’s MSE was higher, reflecting reduced responsiveness to unchanging stimuli due to adaptation (*t*_9_=-4.87*, p*=4.29e^-6^). By contrast, for sequences with periodic motion changes, AdaptNet’s MSE following each change was significantly lower than MotionNet-R, indicating better responsiveness in detecting change (*t*_9_=10.09, *p*=8.45e^-24^). This behavior is consistent with adaptive coding in sensory neurons, which prioritizes novel or changing stimuli to conserve metabolic resources and prevent saturation from constant stimuli (Fairhall et al., 2001). Further, AdaptNet showed significantly lower MT unit activity in both conditions (change: *t*_9_=159.3 *p=*0; constant: *t_9_*=13.34*, p=*9.07e^-24^; Fig. 5a, left), consistent with the hypothesis that adaptation enhances metabolic efficiency by reducing neuronal firing in response to prolonged or repeated stimulation (Ganguly et al., 2024; Yi & Grill, 2019).; constant: *t*_9_=*13.34, p*=9.07e^-24^; Fig. 5a, left). This lower activity supports the hypothesis that adaptation enhances metabolic efficiency by reducing neuronal firing in response to prolonged or repeated stimulation (Ganguly et al., 2024; Yi & Grill, 2019). This reflects a real biological trade-off: adaptation conserves energy (lower firing over time) but can reduce fidelity for sustained stimuli. In nature, this is often acceptable because events that change are typically considered more ecologically relevant than those that remain the same (Tesileanu et al., 2022).

**Figure 5.**
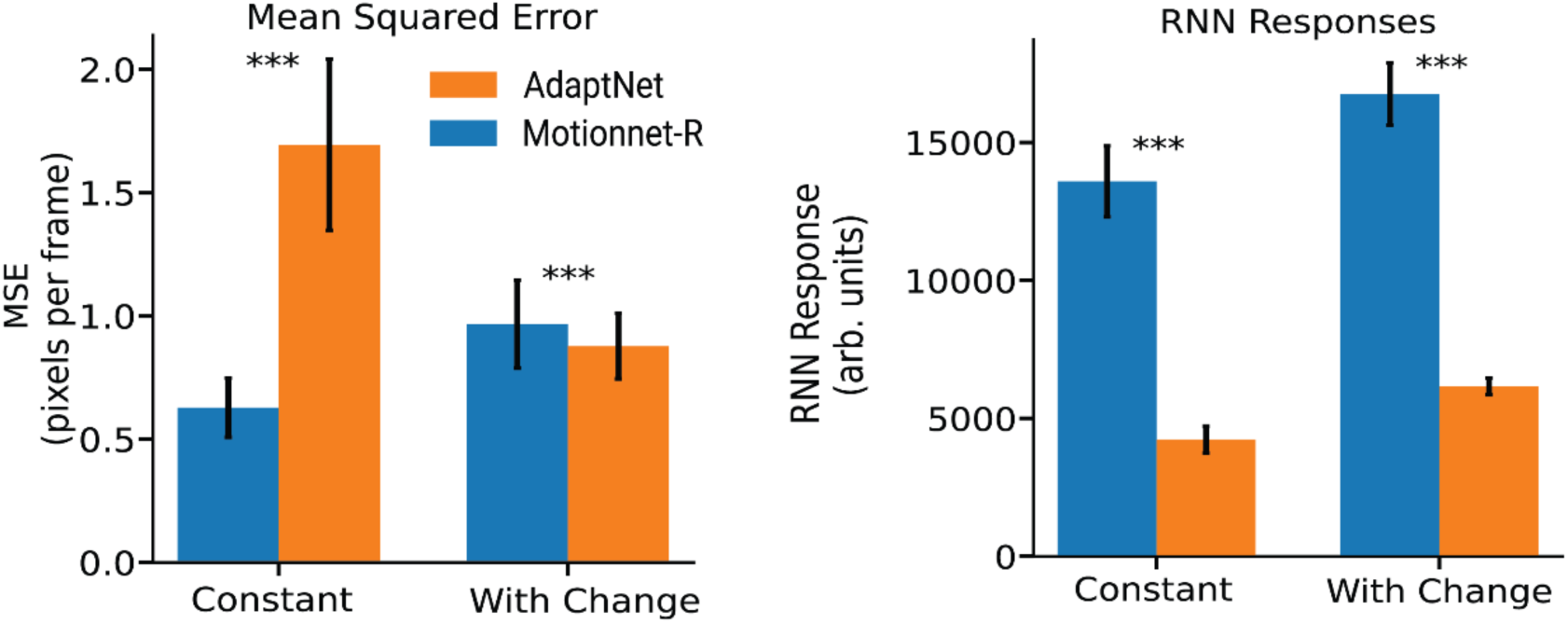
Adaptation improves sensitivity to change and increases efficiency. **a)** The mean squared error (MSE, left) and total MT layer activity (right) for MotionNet-R and AdaptNet for image sequences of constant motion (orange) and periodic motion changes (blue). Error bars indicate ±SD across multiple networks, and asterisks denote statistical significance (p<.001).

To further investigate the networks’ sensitivity to constant and changing motion, we tested both with a dot-motion stimulus that moved in one velocity for nine frames, and then changed either remained the same or changed to different velocity on the tenth frame. We recorded the direction of the networks’ estimate for the final frame and compared this to the actual (ground truth) direction of the final velocity. Overall, we found that AdaptNet’s estimates were closer to the actual velocity than those of MotionNet-R (Fig 6a).

**Figure 6.**
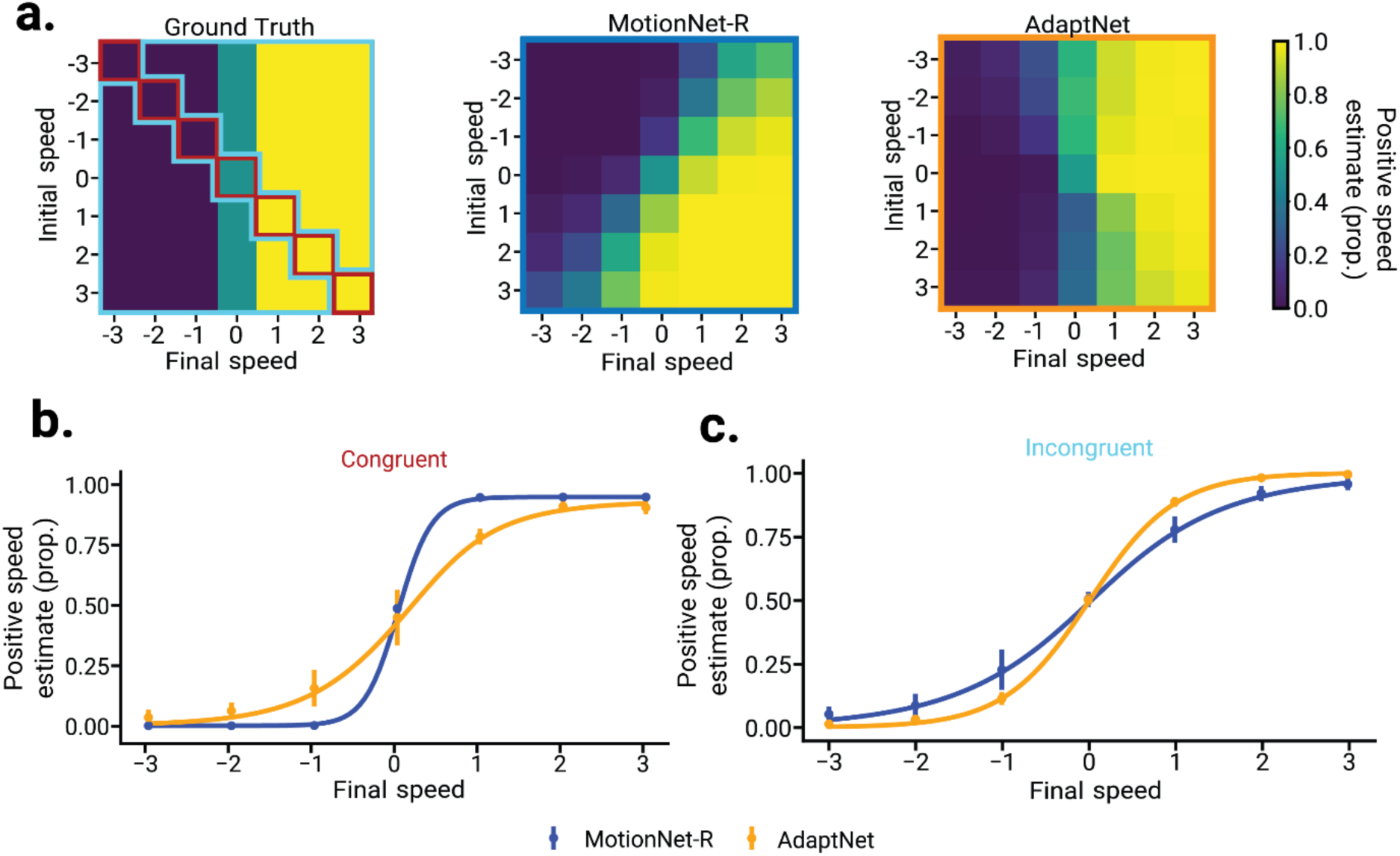
Adaptation enhances sensitivity to change. **a)** The proportion of positive velocity estimates along the (combined x and y) tested dimensions for each combination of initial (first nine frames) and final (tenth) frame speeds, for MotionNet-R (center) and AdaptNet (right); the ground truth (left) is displayed for reference. **b)** The same as **(a,** MotionNet-R & AdaptNet), but just for the (congruent) combinations of speeds along the diagonal **(a,** Ground truth, red bordered region). **c)** The same as **(a,** MotionNet-R & AdaptNet), but averaged across the remaining (incongruent) combinations of speeds **(a,** Ground truth, cyan bordered region). Sigmoid functions were fit to the data for both networks, and the steeper slopes indicate higher sensitivity. Error bars in (**b** & **c**) indicate ±SD across multiple networks.

We compared the sensitivity of each network to congruent and incongruent motion sequences by fitting their performance across final speeds with a sigmoid function (Fig. 6b**-c**). For the congruent condition, where the speed was constant across all frames, we found that MotionNet-R was more sensitive to speed (as indicated by the steeper slope; Fig. 6b). This is expected, because AdaptNet’s accuracy reduces in response to constant stimuli. By contrast, for incongruent sequences, where the final speed changed, we found that AdaptNet was more sensitive (Fig. 6c). This increased sensitivity to change captures captures what is seen in biology across neural systems in different species, where adaptation is thought to amplify sensitivity to changes in the environment (Fairhall et al., 2001; Luczak & Kubo, 2022).

Neurons rapidly reset their adaptation when newly activated, suppressing previously adapted neurons, enhancing responsiveness to motion changes. Such improvement in response sensitivity supports the idea that incorporating adaptive mechanisms could introduce functional benefits in a neural network. Improved sensitivity to dynamic changes and increased processing efficiency have also previously been theorized to be the functional advantages of adaptation in biological systems. By capturing these key aspects of neural adaptation, AdaptNet offers a valuable model for exploring how adaptation shapes motion perception and neural computation.

## DISCUSSION

Here we trained two artificial neural networks, MotionNet-R and AdaptNet, to estimate motion from natural image sequences. Our primary aim was to investigate whether introducing adaptive mechanisms into motion processing networks would recapitulate phenomena associated with motion adaptation in biological neural networks and perception. Our results show that AdaptNet, but not MotionNet-R, exhibited the well-known perceptual phenomenon associated with motion adaptation, the MAE, and further provided a biologically explicit explanation for the effect. We then showed that AdaptNet was both more energy efficient and sensitive to changes in stimulus motion compared to the non-adaptive network (MotionNet-R).

The relevance of insights gleaned from ANNs into how biological neural networks operate is predicated on the correspondence between their functions. Thus, we began by validating the emergent properties of the ANNs with previously observed aspects of biological motion processing. We found similar hierarchical motion processing in the ANNs as observed in the visual cortex. In particular, the V1 units of both networks developed response properties that aligned with those of V1 neurons in macaque, such as the preference of V1 neurons for relatively lower speeds compared to MT neurons (DeAngelis et al., 1999; Hubel & Wiesel, 1962). The response of the units to plaid stimuli also varied considerably; while V1 units responded to individual components of the plaid grating, MT units responded to the pattern motion of the stimulus. This further validates the models’ biological plausibility (Rideaux & Welchman, 2020; Zaharia et al., 2019) and reflects the role of MT in integrating local motion signals into a global percept (Maunsell & Van Essen, 1983; Perrone & Thiele, 2001).

We next tested our networks with stimuli that ceased moving after a period of constant motion, to compare their responses to those observed in psychophysical studies of motion adaptation (Mather & Harris, 1998). The adaptive network (AdaptNet) overshot the stationary (zero) motion estimate when the motion ceased, providing an estimate of motion in the opposite direction. This captures what is observed during psychophysical accounts of the phenomenon, suggesting that our network recapitulates a key aspect of adaptation. We then showed that the MAE in AdaptNet occurred because of a temporary reduction in the baseline activity of MT units, induced by prolonged exposure to a single motion direction, which perturbed the otherwise balanced mutual suppression between units tuned to different directions that exists during exposure to static images. This mechanism has previously been hypothesized as the basis of the MAE (Allman et al., 1985; Barlow & Hill, 1963), and our findings provide empirical support for this account.

On analyzing the MT unit’s responses during MAE, we also observed that AdaptNet showed a decreased latency in response to the onset of motion, compared to MotionNet-R. We investigated this further by measuring the number of frames required by each network to reach their peak output response following a change in the input motion and found that AdaptNet reached its peak 2-3 frames earlier than MotionNet-R. In particular, AdaptNet’s response seemed to peak upon motion onset while MotionNet-R’s response gradually increased over time. The former is consistent with rapid processing of changing input while the latter is more suited to integration of information over time. Our results may indicate that the degree to which biological neurons adapt may be related to their role as change detectors or temporal integrators.

Adaptation is thought to support healthy brain function by reducing the metabolic expenditure of processing sensory information (Kohn, 2007; Niven, 2016). Consistent with this, we found that AdaptNet used less energy to represent motion, both in V1 and MT, operationalized as reduced absolute unit activity within each layer. Efficiency is crucial in biological systems, where rapid and accurate sensory processing must be balanced with metabolic constraints (Lennie, 2003). The increased energy efficiency in AdaptNet came at the cost of reduced estimation accuracy for prolonged constant motion. However, we found that AdaptNet produced more accurate estimates in response to changes in motion. This improved accuracy was mirrored by a corresponding increase in sensitivity. These emergent properties support the notion that adaptation supports perception by increasing sensitivity to changes in sensory input (Fairhall et al., 2001; Rideaux et al., 2023; Wark et al., 2007).

This study addresses key gaps in the field of computational neuroscience. Previous models such as those by Simoncelli and Heeger (1998) and Rust et al. (2006), accounted for direction and speed tuning but did not incorporate adaptation. Accounting for this mechanism enabled us to provide a more realistic simulation that not only replicates the known tuning properties of motion processing neurons, but also captures the dynamic aspects of neural adaptation (Clifford & Wenderoth, 1999; Krekelberg et al., 2006).

The current study provides further insight into well-known perceptual phenomena (MAE), supports existing ecological explanations for adaptation, and reveals a new potential benefit of this canonical neural feature (reduced latency). These findings extend the current understanding of motion processing and sensory adaptation, but they may also have important implications for artificial neural systems. That is, the increased sensitivity and reduced latency to motion changes in AdaptNet highlights the potential for adaptation mechanisms to improve the performance of artificial vision systems, particularly in dynamic environments where rapid detection of motion changes is desirable (Cox & Dean, 2014; Tan & van Dijken, 2023).

### Limitations

We acknowledge some limitations of our study that should be considered. While AdaptNet successfully captured certain aspects of neural adaptation, the adaptation in our network was operationalized with a relatively simple mechanism. By design, this simplifies the complexity of adaptation in biological systems, such as the influence of neurotransmitters, ion channel dynamics, and modulatory inputs from other brain regions (Benda & Herz, 2003). Our study was conducted on a dataset of natural image sequences from the Berkeley Segmentation Dataset (Martin et al., 2001). While this provides a relatively naturalistic visual diet for our artificial systems, it does not capture all the variability present in real-world visual environments.

We did not compare the performance of AdaptNet with other existing motion processing networks becuase our networks were not designed to compete against computational benchmarks, rather, we sought to follow a biologically plausible training regime while constraining the network’s size and complexity, in order to interrogate the properties that emerge. Further, we sought to study the isolated effects of adaptation. To this end we used MotionNet-R as a baseline network, as previous studies have validated the motion processing properties of similar architectures (Rideaux & Welchman, 2020, 2021). This strategy allowed us to isolate the effect of adding adaptation to a relatively well-understood architecture.

### Future Work

Future work could increase the biological plausibility of the network by using more realistic spiking models of neurons (Izhikevich, 2003). Extending the networks’ input space to include more complex and varied stimuli like 3D motion could highlight other domains where adaptation influences sensory processing and perception (Idrees et al., 2024; Latimer et al., 2019; Rideaux et al., 2021).

With this study, we sought to demonstrate the role of adaptation mechanisms in motion processing systems. Incorporating adaptation into a motion processing network enabled us to replicate key biological phenomena that have been observed in the dorsal stream of the visual system. Further to this, we observed a higher energy efficiency and enhanced sensitivity to varying stimuli in a neural network with adaptation. These findings offer novel insights on the role of adaptation in motion processing and neural processing more broadly.

## Acknowledgements

This work was supported by Australian Research Council (ARC) Discovery Early Career Researcher Awards awarded to RR (DE210100790) a National Health and Medical Research Council (NHMRC; Australia) Investigator Grant (2026318).

## Supplementary Figures

**Supplementary Figure 1.**
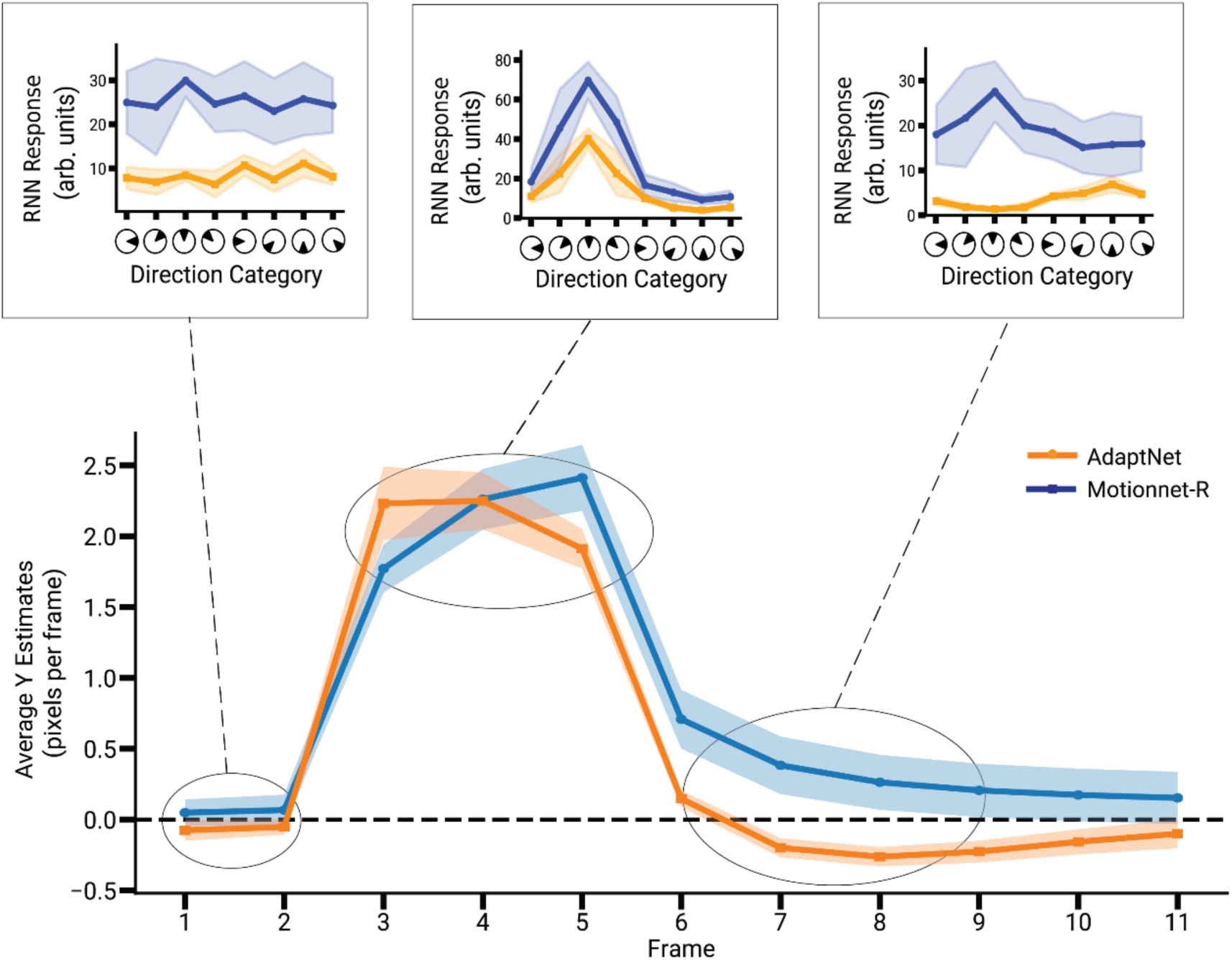
Motion Aftereffect in AdaptNet (Y component). Time course of the networks’ Y responses to an image sequence consisting of a stationary stage (2 frames), followed by a moving stage (3 frames), and then a return to stationary (5 frames). The lower plot shows MotionNet-R and AdaptNet’s motion estimation over time. The upper insets (showing RNN responses during frames 2, 4 and 8 representing the stationary, moving and final phases of the sequence respectively) show the average activity of MT units, binned according to their directional tuning, during different stages of the sequence. Shaded regions indicate ±SD across multiple networks.

**Supplementary Figure 2.**
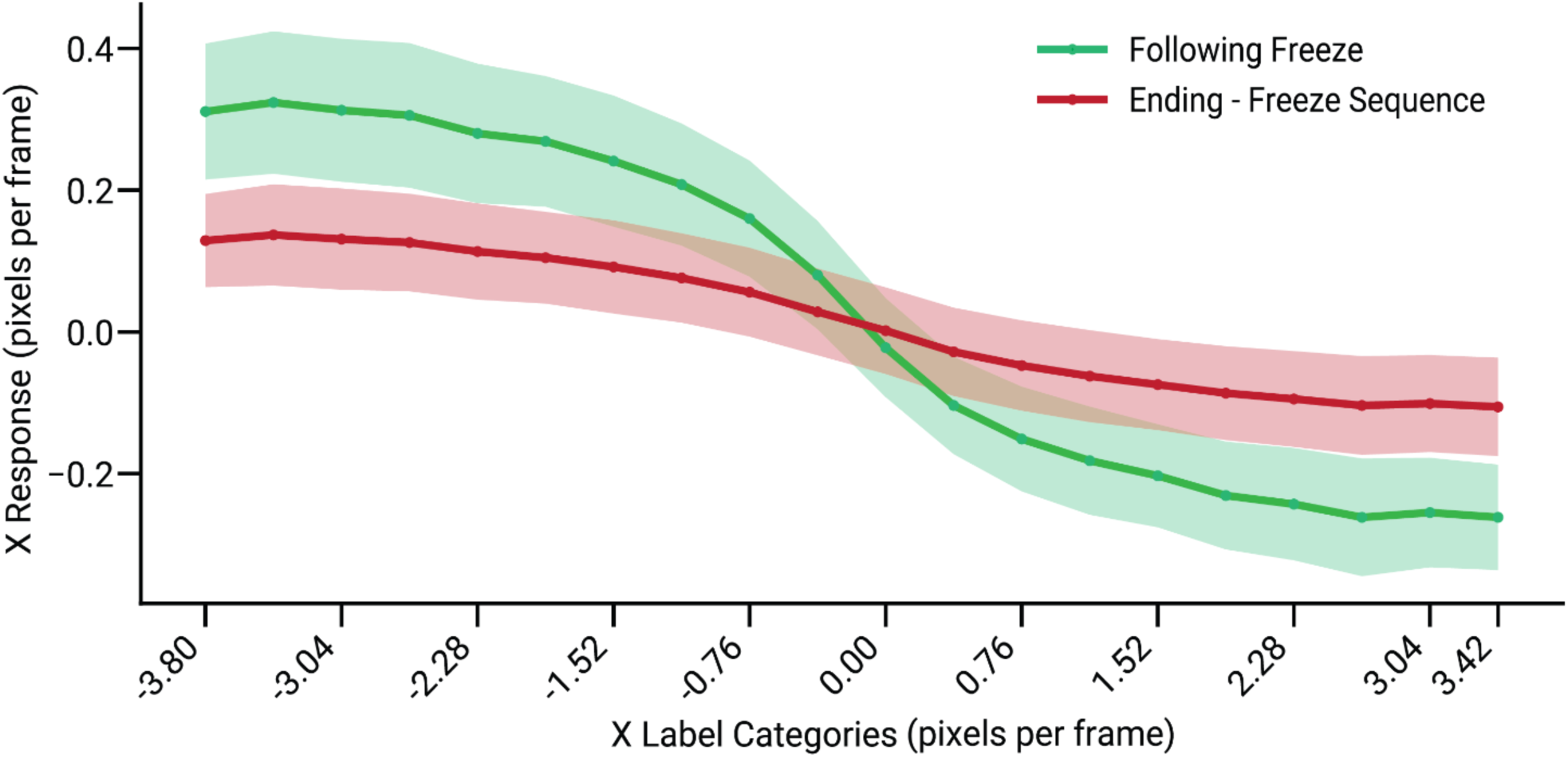
Motion Aftereffect observed across a range of sequence velocities. Response magnitudes of the X component of AdaptNet are plotted immediately after the motion sequence has been stopped (green) along with the response magnitudes of AdaptNet towards the end of the sequence (red). The graphs show how across a variety of input sequence velocities, AdaptNet first overshoots to make the opposite prediction and then corrects itself towards the end. Shaded regions indicate ±SD across multiple networks.

**Supplementary Figure 3.**
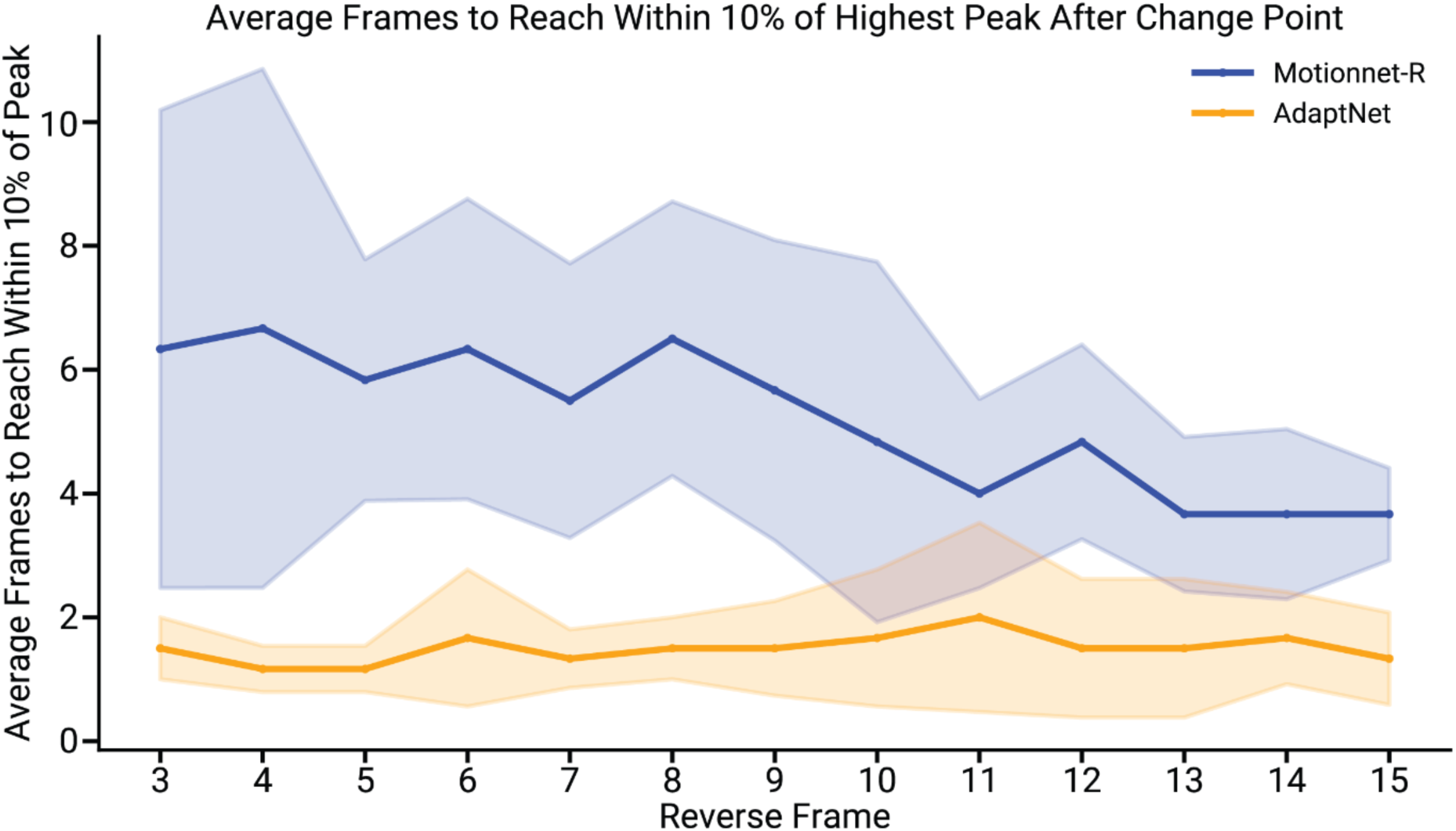
AdaptNet shows lower response latency. Graphs showing the average number of frames taken by the networks to reach within 10% of their maximum response value, following a change in the velocity of the input sequence. AdaptNet reaches this range within 1-2 frames on average regardless of how adapted the state of the network is, just before the change in velocity while MotionNet-R trails behind in terms of its response. Shaded regions indicate ±SD across multiple networks.

## REFERENCES

Albright, T. D. (1984). Direction and orientation selectivity of neurons in visual area MT of the macaque. Journal of Neurophysiology, 52(6), 1106–1130. 10.1152/jn.1984.52.6.1106

Allman, J., Miezin, F., & McGuinness, E. (1985). Direction- and Velocity-Specific Responses from beyond the Classical Receptive Field in the Middle Temporal Visual Area (MT). Perception, 14(2), 105–126. 10.1068/p140105

Bair, W., & Movshon, J. A. (2004). Adaptive Temporal Integration of Motion in Direction-Selective Neurons in Macaque Visual Cortex. The Journal of Neuroscience, 24(33), Article 33. 10.1523/JNEUROSCI.0554-04.2004

Barlow, H. B., & Hill, R. M. (1963). Evidence for a Physiological Explanation of the Waterfall Phenomenon and Figural After-effects. Nature, 200(4913), 1345–1347. 10.1038/2001345a0

Barlow, HB., & Foldiak, P. (1989). Adaptation and decorrelation in the cortex. In C. Miall, RM. Durbin, & GJ. Mitchison (Eds.), The Computing Neuron (pp. 54–72). Addison Wesley.

Benda, J., & Herz, A. V. M. (2003). A universal model for spike-frequency adaptation. Neural Computation, 15(11), 2523–2564. 10.1162/089976603322385063

Born, R. T., & Bradley, D. C. (2005). STRUCTURE AND FUNCTION OF VISUAL AREA MT. Annual Review of Neuroscience, 28(1), Article 1. 10.1146/annurev.neuro.26.041002.131052

Britten, K. H. (2008). Mechanisms of self-motion perception. Annual Review of Neuroscience, 31, 389–410. 10.1146/annurev.neuro.29.051605.112953

Burr, D., & Thompson, P. (2011). Motion psychophysics: 1985–2010. Vision Research, 51(13), Article 13. 10.1016/j.visres.2011.02.008

Cadieu CF, Hong H, Yamins DL, Pinto N, Ardila D, Solomon EA, Majaj NJ, DiCarlo JJ. Deep neural networks rival the representation of primate IT cortex for core visual object recognition. PLoS Comput Biol. 2014 Dec 18;10(12):e1003963. doi: 10.1371/journal.pcbi.1003963. PMID: 25521294; PMCID: PMC4270441.

Carandini, M., Heeger, D. J., & Anthony Movshon, J. (1999). Linearity and Gain Control in V1 Simple Cells. In P. S. Ulinski, E. G. Jones, & A. Peters (Eds.), Cerebral Cortex (Vol. 13, pp. 401–443). Springer US. 10.1007/978-1-4615-4903-1_7

Churchland, M. M., & Lisberger, S. G. (2001). Shifts in the population response in the middle temporal visual area parallel perceptual and motor illusions produced by apparent motion. The Journal of Neuroscience, 21(23), 9387–9402.

Clifford CW, Webster MA, Stanley GB, Stocker AA, Kohn A, Sharpee TO, Schwartz O. Visual adaptation: neural, psychological and computational aspects. Vision Res. 2007 Nov;47(25):3125–31. doi: 10.1016/j.visres.2007.08.023. Epub 2007 Oct 22. PMID: 17936871.

Clifford, C. W. G., & Wenderoth, P. (1999). Adaptation to temporal modulation can enhance differential speed sensitivity. Vision Research, 39(26), Article 26. 10.1016/S0042-6989(99)00151-0

Cox, D. D., & Dean, T. (2014). Neural Networks and Neuroscience-Inspired Computer Vision. Current Biology, 24(18), R921–R929. 10.1016/j.cub.2014.08.026

Cuturi, L. F., & MacNeilage, P. R. (2014). Optic flow induces nonvisual self-motion aftereffects. Current Biology: CB, 24(23), 2817–2821. 10.1016/j.cub.2014.10.015

De Valois, R. L., Albrecht, D. G., & Thorell, L. G. (1982). Spatial frequency selectivity of cells in macaque visual cortex. Vision Research, 22(5), Article 5. 10.1016/0042-6989(82)90113-4

DeAngelis, G. C., Ghose, G. M., Ohzawa, I., & Freeman, R. D. (1999). Functional Micro-Organization of Primary Visual Cortex: Receptive Field Analysis of Nearby Neurons. Journal of Neuroscience, 19(10), 4046–4064. 10.1523/JNEUROSCI.19-10-04046.1999

Duong, L. R., Bredenberg, C., Heeger, D. J., & Simoncelli, E. P. (2023). *Adaptive coding efficiency in recurrent cortical circuits via gain control* (No. arXiv:2305.19869). arXiv. 10.48550/arXiv.2305.19869

Edwards, M., & Rideaux, R. (2013). How many motion signals can be simultaneously perceived? Vision Research, 76, 11–16. 10.1016/j.visres.2012.10.004

Fairhall AL, Lewen GD, Bialek W, de Ruyter Van Steveninck RR. Efficiency and ambiguity in an adaptive neural code. Nature. 2001 Aug;412(6849):787–792. DOI: 10.1038/35090500. PMID: 11518957.

Ganguly, C., Bezugam, S. S., Abs, E., Payvand, M., Dey, S., & Suri, M. (2024). Spike frequency adaptation: Bridging neural models and neuromorphic applications. Communications Engineering, 3(1), 1–13. 10.1038/s44172-024-00165-9

Glasser, D. M., Tsui, J. M. G., Pack, C. C., & Tadin, D. (2011). Perceptual and neural consequences of rapid motion adaptation. Proceedings of the National Academy of Sciences, 108(45), Article 45. 10.1073/pnas.1101141108

Grunewald, A., & Lankheet, M. J. M. (1996). The Orthogonal Motion Aftereffect. Perception, 25(1_suppl), 65–65. 10.1068/v96l0805

Guclu, U., & Van Gerven, M. A. J. (2015). Deep Neural Networks Reveal a Gradient in the Complexity of Neural Representations across the Ventral Stream. Journal of Neuroscience, 35(27), Article 27. 10.1523/JNEUROSCI.5023-14.2015

Gundavarapu, A., Chakravarthy, V. S., & Soman, K. (2019). A Model of Motion Processing in the Visual Cortex Using Neural Field With Asymmetric Hebbian Learning. Frontiers in Neuroscience, 13, 67. 10.3389/fnins.2019.00067

Heeger, D. J., Boynton, G. M., Demb, J. B., Seidemann, E., & Newsome, W. T. (1999). Motion Opponency in Visual Cortex. The Journal of Neuroscience, 19(16), 7162. 10.1523/JNEUROSCI.19-16-07162.1999

Hietanen, M. A., Crowder, N. A., Price, N. S. C., & Ibbotson, M. R. (2007). Influence of adapting speed on speed and contrast coding in the primary visual cortex of the cat. The Journal of Physiology, 584(Pt 2), 451–462. 10.1113/jphysiol.2007.131631

Hu, B., Garrett, M. E., Groblewski, P. A., Ollerenshaw, D. R., Shang, J., Roll, K., Manavi, S., Koch, C., Olsen, S. R., & Mihalas, S. (2021). Adaptation supports short-term memory in a visual change detection task. PLOS Computational Biology, 17(9), e1009246. 10.1371/journal.pcbi.1009246

Hubel DH, Wiesel TN. Receptive fields, binocular interaction and functional architecture in the cat’s visual cortex. J Physiol. 1962 Jan;160(1):106–54. doi: 10.1113/jphysiol.1962.sp006837. PMID: 14449617; PMCID: PMC1359523.

Idrees, S., Manookin, M. B., Rieke, F., Field, G. D., & Zylberberg, J. (2024). Biophysical neural adaptation mechanisms enable artificial neural networks to capture dynamic retinal computation. Nature Communications, 15(1), 5957. 10.1038/s41467-024-50114-5

Izhikevich, E. M. (2003). Simple model of spiking neurons. IEEE Transactions on Neural Networks, 14(6), 1569–1572. IEEE Transactions on Neural Networks. 10.1109/TNN.2003.820440

Johnston, A., Arnold, D. H., & Nishida, S. (2006). Spatially localized distortions of event time. Current Biology: CB, 16(5), 472–479. 10.1016/j.cub.2006.01.032

Kar, K., Kubilius, J., Schmidt, K., Issa, E. B., & DiCarlo, J. J. (2019). Evidence that recurrent circuits are critical to the ventral stream’s execution of core object recognition behavior. Nature Neuroscience, 22(6), 974–983. 10.1038/s41593-019-0392-5

Khaligh-Razavi SM, Kriegeskorte N. Deep supervised, but not unsupervised, models may explain IT cortical representation. PLoS Comput Biol. 2014 Nov 6;10(11):e1003915. doi: 10.1371/journal.pcbi.1003915. PMID: 25375136; PMCID: PMC4222664.

Kingma, D. P., & Ba, J. (2017). Adam: A Method for Stochastic Optimization (No. arXiv:1412.6980). arXiv. 10.48550/arXiv.1412.6980

Kohn, A. (2007). Visual Adaptation: Physiology, Mechanisms, and Functional Benefits. Journal of Neurophysiology, 97(5), 3155–3164. 10.1152/jn.00086.2007

Kohn, A., & Movshon, J. A. (2003). Neuronal adaptation to visual motion in area MT of the macaque. Neuron, 39(4), 681–691. 10.1016/s0896-6273(03)00438-0

Kohn, A., & Movshon, J. A. (2004). Adaptation changes the direction tuning of macaque MT neurons. Nature Neuroscience, 7(7), Article 7. 10.1038/nn1267

Krekelberg, B., Boynton, G. M., & Van Wezel, R. J. A. (2006). Adaptation: From single cells to BOLD signals. Trends in Neurosciences, 29(5), Article 5. 10.1016/j.tins.2006.02.008

Kriegeskorte, N., & Douglas, P. K. (2018). Cognitive computational neuroscience. Nature Neuroscience, 21(9), Article 9. 10.1038/s41593-018-0210-5

Latimer, K. W., Barbera, D., Sokoletsky, M., Awwad, B., Katz, Y., Nelken, I., Lampl, I., Fairhall, A. L., & Priebe, N. J. (2019). Multiple Timescales Account for Adaptive Responses across Sensory Cortices. Journal of Neuroscience, 39(50), 10019– 10033. 10.1523/JNEUROSCI.1642-19.2019

Lee, A. L. F. (2018). The contribution of local and global motion adaptation in the repulsive direction aftereffect. Journal of Vision, 18(12), 2. 10.1167/18.12.2

Lennie, P. (2003). The Cost of Cortical Computation. Current Biology, 13(6), 493–497. 10.1016/S0960-9822(03)00135-0

Li, L., Ji, X., Liang, F., Li, Y., Xiao, Z., Tao, H. W., & Zhang, L. I. (2014). A Feedforward Inhibitory Circuit Mediates Lateral Refinement of Sensory Representation in Upper Layer 2/3 of Mouse Primary Auditory Cortex. Journal of Neuroscience, 34(41), 13670–13683. 10.1523/JNEUROSCI.1516-14.2014

Luczak, A., & Kubo, Y. (2022). Predictive Neuronal Adaptation as a Basis for Consciousness. Frontiers in Systems Neuroscience, 15. 10.3389/fnsys.2021.767461

Mao, J., Rothkopf, C. A., & Stocker, A. A. (2025). Adaptation optimizes sensory encoding for future stimuli. PLOS Computational Biology, 21(1), e1012746. 10.1371/journal.pcbi.1012746

Martin, D., Fowlkes, C., Tal, D., & Malik, J. (2001). A database of human segmented natural images and its application to evaluating segmentation algorithms and measuring ecological statistics. Proceedings Eighth IEEE International Conference on Computer Vision. ICCV 2001, *2*, 416–423 vol.2. 10.1109/ICCV.2001.937655

Mather, G., & Harris, J. (1998). Theoretical Models of the Motion Aftereffect. In G. Mather, F. Verstraten, & S. Anstis (Eds.), The Motion Aftereffect (pp. 153–181). The MIT Press. 10.7551/mitpress/4779.003.0008

Mather, G., Verstraten, F., & Anstis, S. (Eds.). (1998). The Motion Aftereffect: A Modern Perspective. The MIT Press. 10.7551/mitpress/4779.001.0001

Maunsell, J., & Van Essen, D. (1983). The connections of the middle temporal visual area (MT) and their relationship to a cortical hierarchy in the macaque monkey. The Journal of Neuroscience, 3(12), Article 12. 10.1523/JNEUROSCI.03-12-02563.1983

Movshon, J. A., & Newsome, W. T. (1996). Visual Response Properties of Striate Cortical Neurons Projecting to Area MT in Macaque Monkeys. The Journal of Neuroscience, 16(23), Article 23. 10.1523/JNEUROSCI.16-23-07733.1996

Movshon, J., Adelson, E. H., Gizzi, M. S., & Newsome, W. T. (1985a). The analysis of moving visual patterns. In C. Chagas, R. Gattass, & C. Gross (Eds.), Pattern recognition mechanisms (pp. 117–151). Vatican Press.

Movshon, J., Adelson, E. H., Gizzi, M. S., & Newsome, W. T. (1985b). The analysis of moving visual patterns. In C. Chagas, R. Gattass, & C. Gross (Eds.), Pattern recognition mechanisms (pp. 117–151). Vatican Press.

Nishida, S. (2011). Advancement of motion psychophysics: Review 2001-2010. Journal of Vision, 11(5), Article 5. 10.1167/11.5.11

Niven, J. E. (2016). Neuronal energy consumption: Biophysics, efficiency and evolution. Current Opinion in Neurobiology, 41, 129–135. 10.1016/j.conb.2016.09.004

Patterson, C. A., Duijnhouwer, J., Wissig, S. C., Krekelberg, B., & Kohn, A. (2014). Similar adaptation effects in primary visual cortex and area MT of the macaque monkey under matched stimulus conditions. Journal of Neurophysiology, 111(6), 1203–1213. 10.1152/jn.00030.2013

Patterson CA, Wissig SC, Kohn A. Distinct effects of brief and prolonged adaptation on orientation tuning in primary visual cortex. J Neurosci. 2013 Jan 9;33(2):532–43. doi: 10.1523/JNEUROSCI.3345-12.2013. PMID: 23303933; PMCID: PMC3710132.

Perrone, J. A., & Thiele, A. (2001). Speed skills: Measuring the visual speed analyzing properties of primate MT neurons. Nature Neuroscience, 4(5), 526–532. 10.1038/87480

Priebe, N. J., Churchland, M. M., & Lisberger, S. G. (2002). Constraints on the Source of Short-Term Motion Adaptation in Macaque Area MT. I. The Role of Input and Intrinsic Mechanisms. Journal of Neurophysiology, 88(1), 354–369. 10.1152/jn.00852.2001

Priebe, N. J., Lisberger, S. G., & Movshon, J. A. (2006). Tuning for Spatiotemporal Frequency and Speed in Directionally Selective Neurons of Macaque Striate Cortex. Journal of Neuroscience, 26(11), 2941–2950. 10.1523/JNEUROSCI.3936-05.2006

Qian, N., & Andersen, R. (1994). Transparent motion perception as detection of unbalanced motion signals. II. Physiology. The Journal of Neuroscience, 14(12), Article 12. 10.1523/JNEUROSCI.14-12-07367.1994

Qiao, L., & Shen, Q. (2021). Human Action Recognition Technology in Dance Video Image. Scientific Programming, 2021(1), 6144762. 10.1155/2021/6144762

Rideaux, R., & Edwards, M. (2014). Information extraction during simultaneous motion processing. Vision Research, 95, 1–10. 10.1016/j.visres.2013.11.007

Rideaux, R., & Harrison, W. J. (2019). Border ownership-dependent tilt aftereffect for shape defined by binocular disparity and motion parallax. Journal of Neurophysiology, 121(5), 1917. 10.1152/jn.00111.2019

Rideaux, R., Storrs, K. R., Maiello, G., & Welchman, A. E. (2021). How multisensory neurons solve causal inference. Proceedings of the National Academy of Sciences of the United States of America, 118(32), e2106235118. 10.1073/pnas.2106235118

Rideaux, R., & Welchman, A. E. (2020). But Still It Moves: Static Image Statistics Underlie How We See Motion. Journal of Neuroscience, 40(12), 2538–2552. 10.1523/JNEUROSCI.2760-19.2020

Rideaux, R., & Welchman, A. E. (2021). Exploring and explaining properties of motion processing in biological brains using a neural network. Journal of Vision, 21(2), 11. 10.1167/jov.21.2.11

Rideaux R, West RK, Rangelov D, Mattingley JB. Distinct early and late neural mechanisms regulate feature-specific sensory adaptation in the human visual system. Proc Natl Acad Sci U S A. 2023 Feb 7;120(6):e2216192120. doi: 10.1073/pnas.2216192120. Epub 2023 Feb 1. PMID: 36724257; PMCID: PMC9963156.

Rust, N. C., Mante, V., Simoncelli, E. P., & Movshon, J. A. (2006). How MT cells analyze the motion of visual patterns. Nature Neuroscience, 9(11), Article 11. 10.1038/nn1786

Sakano, Y., & Allison, R. S. (2014). Aftereffect of motion-in-depth based on binocular cues: Effects of adaptation duration, interocular correlation, and temporal correlation. Journal of Vision, 14(8), 21. 10.1167/14.8.21

Simoncelli, E. P., & Olshausen, B. A. (2001). Natural Image Statistics and Neural Representation. Annual Review of Neuroscience, 24(1), Article 1. 10.1146/annurev.neuro.24.1.1193

Snowden, R. J., Treue, S., & Andersen, R. A. (1992). The response of neurons in areas V1 and MT of the alert rhesus monkey to moving random dot patterns. Experimental Brain Research, 88(2), Article 2. 10.1007/BF02259114

Solomon, S. G., & Kohn, A. (2014). Moving Sensory Adaptation beyond Suppressive Effects in Single Neurons. Current Biology, 24(20), R1012–R1022. 10.1016/j.cub.2014.09.001

Srivastava, N., Hinton, G., Krizhevsky, A., Sutskever, I., & Salakhutdinov, R. (2014). Dropout: A Simple Way to Prevent Neural Networks from Overfitting. Journal of Machine Learning Research, 15(56), 1929–1958.

Tan, A. Y. Y., Brown, B. D., Scholl, B., Mohanty, D., & Priebe, N. J. (2011). Orientation Selectivity of Synaptic Input to Neurons in Mouse and Cat Primary Visual Cortex. Journal of Neuroscience, 31(34), 12339–12350. 10.1523/JNEUROSCI.2039-11.2011

Tan, H., & van Dijken, S. (2023). Dynamic machine vision with retinomorphic photomemristor-reservoir computing. Nature Communications, 14(1), 2169. 10.1038/s41467-023-37886-y

Tesileanu, T., Piasini, E., & Balasubramanian, V. (2022). Efficient processing of natural scenes in visual cortex. Frontiers in Cellular Neuroscience, 16. 10.3389/fncel.2022.1006703

Theusner, S., De Lussanet, M. H. E., & Lappe, M. (2011). Adaptation to biological motion leads to a motion and a form aftereffect. *Attention, Perception*, & Psychophysics, 73(6), Article 6. 10.3758/s13414-011-0133-7

Wainwright, M. J. (1999). Visual adaptation as optimal information transmission. Vision Research, 39(23), Article 23. 10.1016/S0042-6989(99)00101-7

Wark, B., Lundstrom, B. N., & Fairhall, A. (2007). Sensory adaptation. Current Opinion in Neurobiology, 17(4), 423–429. 10.1016/j.conb.2007.07.001

Wissig, S. C., & Kohn, A. (2012). The influence of surround suppression on adaptation effects in primary visual cortex. Journal of Neurophysiology, 107(12), 3370–3384. 10.1152/jn.00739.2011

Yamins DL, DiCarlo JJ. Using goal-driven deep learning models to understand sensory cortex. Nat Neurosci. 2016 Mar;19(3):356–65. doi: 10.1038/nn.4244. PMID: 26906502.

Yamins, D. L. K., Hong, H., Cadieu, C. F., Solomon, E. A., Seibert, D., & DiCarlo, J. J. (2014). Performance-optimized hierarchical models predict neural responses in higher visual cortex. Proceedings of the National Academy of Sciences, 111(23), 8619–8624. 10.1073/pnas.1403112111

Yi, G., & Grill, W. M. (2019). Average firing rate rather than temporal pattern determines metabolic cost of activity in thalamocortical relay neurons. Scientific Reports, 9(1), 6940. 10.1038/s41598-019-43460-8

Zaharia, A. D., Goris, R. L. T., Movshon, J. A., & Simoncelli, E. P. (2019). Compound Stimuli Reveal the Structure of Visual Motion Selectivity in Macaque MT Neurons. eNeuro, 6(6). 10.1523/ENEURO.0258-19.2019

Zamboni, E., Kemper, V. G., Goncalves, N. R., Jia, K., Karlaftis, V. M., Bell, S. J., Giorgio, J., Rideaux, R., Goebel, R., & Kourtzi, Z. (2020). Fine-scale computations for adaptive processing in the human brain. eLife, 9, e57637. 10.7554/eLife.57637

Zeiler, M. D., & Fergus, R. (2013). *Visualizing and Understanding Convolutional Networks* (No. arXiv:1311.2901). arXiv. 10.48550/arXiv.1311.2901

Zhou, S., & Yu, Y. (2018). Synaptic Excitatory-Inhibitory Balance Underlying Efficient Neural Coding. Advances in Neurobiology, 21, 85–100. 10.1007/978-3-319-94593-4_5

